# SUMOylation Regulates IRF2BP2 Nuclear Localization and Stability

**DOI:** 10.64898/2026.11.09.765419

**Authors:** Renata Ramalho-Oliveira, Matheus A. Rajão, Vilênia T. Souza, Flavia R.G. Carneiro, João P.B. Viola

**Affiliations:** Program of Immunology and Tumor Biology, Brazilian National Cancer Institute (INCA), Rio de Janeiro, 20231-050, RJ, Brazil; Center of Technological Development in Health (CDTS), Oswaldo Cruz Foundation (FIOCRUZ), Rio de Janeiro, 21040-361, RJ, Brazil; Interdisciplinary Laboratory for Medical Research, Oswaldo Cruz Institute (IOC), Oswaldo Cruz Foundation (FIOCRUZ), Rio de Janeiro, 21040-900, RJ, Brazil

**Author notes:** Corresponding author: Programa de Imunologia e Biologia Tumoral; Instituto Nacional de Câncer (INCA); Rua André Cavalcanti, 37; Centro, Rio de Janeiro, RJ 20231-050, Brazil. **Current address:** Department of Cell Biology and Physiology, Washington University School of Medicine, St Louis, 63110, MO, USA.

**Keywords:** IRF2BP2, SUMOylation, ubiquitin, nuclear localization, stability, importin β1

## Abstract

Interferon Regulatory Factor-2 Binding Protein 2 (IRF2BP2) is a transcriptional cofactor implicated in diverse biological processes. Although it functions primarily as a repressor, it can also activate gene expression. However, the mechanisms governing IRF2BP2 regulation remain poorly understood. Here, we demonstrate that IRF2BP2 interacts with UBE2I, the SUMO E2 ligase, and undergoes multi-SUMOylation, which enhances its nuclear localization. We further show that IRF2BP2 nuclear import is importin-β-dependent and requires the functional nuclear localization signal (NLS). IRF2BP2 preferentially accumulates in transcriptionally active euchromatin in both resting and S-phase cells, consistent with its role in transcriptional control. We further reveal that SUMOylation contributes to IRF2BP2 stability. Moreover, IRF2BP2 is also ubiquitinated, predominantly through K6-linked ubiquitin chains, a non-proteolytic modification that aligns with our observation that IRF2BP2 is not degraded by the proteasome. Together, these findings identify SUMOylation and non-proteolytic ubiquitination as key mechanisms regulating IRF2BP2, highlighting a novel SUMO1-NLS-importin-β-dependent pathway that governs its localization.

## INTRODUCTION

Gene expression is tightly regulated by a complex network of factors acting at multiple cellular levels. Protein-protein interactions play a central role in coordinating signaling pathways and connecting extracellular signals to the transcriptional machinery. The Interferon Regulatory Factor-2 Binding Protein (IRF2BP) family comprises four transcriptional cofactors: IRF2BP1, IRF2BP2 (isoforms A and B), and IRF2BPL. Initially, they were characterized as transcriptional coregulators associated with the repression of interferon-responsive genes through interactions with IRF-2 (1). These proteins share two conserved zinc finger domains: an N-terminal C4 and a C-terminal C3HC4 RING finger domain (1). The C4 zinc finger has been implicated in homo- and heteromeric interactions among IRF2BP family members and interaction with partners (2). Although C4-zinc finger motifs are also found in DNA-binding proteins (3), direct DNA binding by IRF2BP2 remains to be conclusively demonstrated, despite evidence supporting its association with chromatin (2, 4, 5). In contrast, the C-terminal RING domain has been implicated in several protein-protein interactions involving IRF2BP2 (6, 7).

IRF2BP2 accumulates several partners identified which implicates IRF2BP2 involvement in a variety of biological processes including cell cycle (8), apoptosis (9), angiogenesis (10), cell differentiation (4, 11), immune response, inflammation and immunodeficiency (7, 12-16), as well as tumorigenesis (15, 17-23). Furthermore, despite primarily being identified as a negative regulator, IRF2BP2 can also positively regulate the expression and function of various genes and proteins. These include *VEGFA*, which is upregulated in skeletal and cardiac muscle cells after ischemia (10); *CTGF* (Connective Tissue Growth Factor) in gastric cancer and normal cells (19); AP-1 (Activator Protein-1, composed of *JUN* and *FOSL2*) in neuroblastoma, with increased ALK expression (22); and *ZBTB16* (Zinc finger and BTB domain-containing protein 16) during megakaryocytic differentiation (11). However, many known IRF2BP2 targets are negatively regulated: IRF-2 (1); p21 and Bax (9); NFAT1 (6); the pro-apoptotic factor FASTKD2 (2); ATF3 (24); the erythroid differentiation gene program (4); and the AP-1 heterodimer ATF7/JDP2 (15).

In immune and inflammatory modulation, Chen and colleagues (8) studied coronary disease and demonstrated that IRF2BP2 activates the anti-inflammatory transcription factor KLF2, which favors the differentiation of M2 anti-inflammatory macrophages. In addition, Cruz and colleagues (17) showed that IRF2BP2 expression in microglia is important to limit inflammation. It was also reported that IRF2BP2 protein favors the recovery from sepsis-induced cardiomyopathy by mediating anti-inflammatory and anti-apoptotic responses (25) and represses IL-1β in AML cells (26). Furthermore, our group also contributed to understanding IRF2BP2’s immune roles. In fact, IRF2BP2 is an NFAT1 partner inhibiting its transcriptional activity (6) and modulates T lymphocyte activation by repressing proliferation and decreasing cytokines’ expression, such as CD25 (12), which is directly regulated by NFAT1 (27), and CD69 (12). Interestingly, IRF2BP2 mutations and gene variations are related to immunodeficiency and autoimmune diseases (14, 16, 28).

The role of IRF2BP2 in carcinogenesis has been addressed as well, showing both anti- and pro-tumoral roles depending on the cellular context. IRF2BP2 suppresses liver cancer progression through the regulation of the Hippo pathway (18) and was reported in PD-L1/PD-1 immune checkpoint regulation. High levels of PD-L1 and lower expression of IRF2BP2 in basal breast cancer cells were observed (29, 30). It was also demonstrated that IRF2BP2 is stabilized by VGLL4 in the Hippo pathway and represses PD-L1 expression favoring anti-tumor immunity (31). On the other hand, IRF2BP2 seems to have an opposite role in cervical cancer cells, where *CDYL*, an epigenetic regulator, inhibited the PD-L1 expression through transcriptionally downregulating IRF2BP2 (32).

The pro-tumoral role of IRF2BP2 has been investigated in several models. IRF2BP2 expression is upregulated in gastric cancer cell lines and tissues, and elevated IRF2BP2 levels are associated with poor overall survival. Conversely, IRF2BP2 knockdown inhibits proliferation, migration, invasion, and epithelial-mesenchymal transition in gastric cancer cells (19). IRF2BP2 also promotes proliferation in neuroblastoma and triple-negative breast cancer cells (20, 22). In addition, IRF2BP2 is upregulated in Acute Myeloid Leukemia (AML) samples and is required to sustain AML cell proliferation (15). Moreover, studies have reported gene fusions involving IRF2BP2 that differ from the classical fusions observed in some cancers, including the IRF2BP2-CDX1 fusion in mesenchymal chondrosarcoma (33, 34) and the IRF2BP2-RARA fusion in acute promyelocytic leukemia (35). These fusions have been identified in patients with more aggressive tumors or resistance to treatment. Finally, IRF2BP2 has also been shown to be critical for the growth and survival of T-cell Acute Lymphoblastic Leukemia (T-ALL) cells (23).

Although many studies have explored IRF2BP2’s roles, its regulation remains poorly understood. It is known that p53 can bind to a site 9 kb upstream of the IRF2BP2 promoter and activate its expression in DNA damage response (9). IRF2BP2 is also post-transcriptionally regulated. At least two miRNAs, miR-155 (36) and miR-101-3p (19), can target IRF2BP2 mRNA and reduce its expression. Moreover, the lncRNA CHROMR, which is induced by viral infection, sequesters IRF2BP2 consequently releasing the transcription of interferon-stimulated genes (37).

At the post-translational level, it was reported that IRF2BP2 can be SUMOylated at Lys566, a conserved site present in zebrafish and humans (5, 38). In the former, SUMOylation contributes to IRF2BP2’s repressive role, and in the latter, transient deSUMOylation in response to EGF treatment in HeLa cells is important to control early transcription in EGFR signaling (38). It was also suggested that IRF2BP2 could be degraded by the ubiquitin-proteasome system because it is ubiquitin-modified, forming K48 chains in MG132-treated cells (31). Moreover, the protein contains a nuclear localization signal (NLS) (6) and a nearby phosphorylation site that favors its nuclear localization(2). Therefore, the molecular mechanism for IRF2BP2 regulation needs to be largely explored.

To address this gap, we investigated the pathways regulating IRF2BP2 function. Using a yeast two-hybrid screen, we identified UBE2I, a SUMO-conjugating enzyme homologous to yeast Ubc9, as a novel interaction partner of IRF2BP2. This finding pointed to a role for the SUMOylation pathway in the IRF2BP2 regulation. The interaction between IRF2BP2 and UBE2I was further confirmed through pull-down assays and FRET analysis. Our data revealed that IRF2BP2 is modified by SUMO1 at three Lysine residues, and this modification promotes its nuclear import via an importin β-dependent mechanism. Furthermore, IRF2BP2 subcellular localization is dynamic across the cell cycle: it is exclusively nuclear during the S phase but can also be detected in the cytoplasm during other phases. Notably, when localized in the nucleus, IRF2BP2 is enriched in euchromatin regions, suggesting its functional involvement in transcriptional regulation. Together, these findings propose a novel regulatory mechanism for IRF2BP2 involving SUMO1 modification and NLS/importin β-mediated nuclear import.

## RESULTS

### IRF2BP2 interacts with UBE2I

Aiming to identify possible partners that could characterize the molecular regulatory mechanism of IRF2BP2 activity, we performed a yeast two-hybrid screening to identify new IRF2BP2-interacting proteins, using full-length IRF2BP2 as bait (**Figure 1A**). Thirty-five percent of the sequenced clones isolated from a total of 10^7^ yeast colonies transformed with a human B-cell cDNA library encoded the UBE2I (Ubiquitin-like binding enzyme 2I) protein, which is the only SUMO E2 ligase described. To confirm the interaction, we initially retransformed the yeast L40 strain containing the full-length IRF2BP2 or the truncated forms IRF2BP2-RING (amino acids 302 to 571) and IRF2BP2-ZINC (amino acids 1 to 484) fused to the LexA DNA binding domain with full-length UBE2I fused to Gal4p activation domain and analyzed the reporter genes (**Figure 1A**).

**Figure 1.**
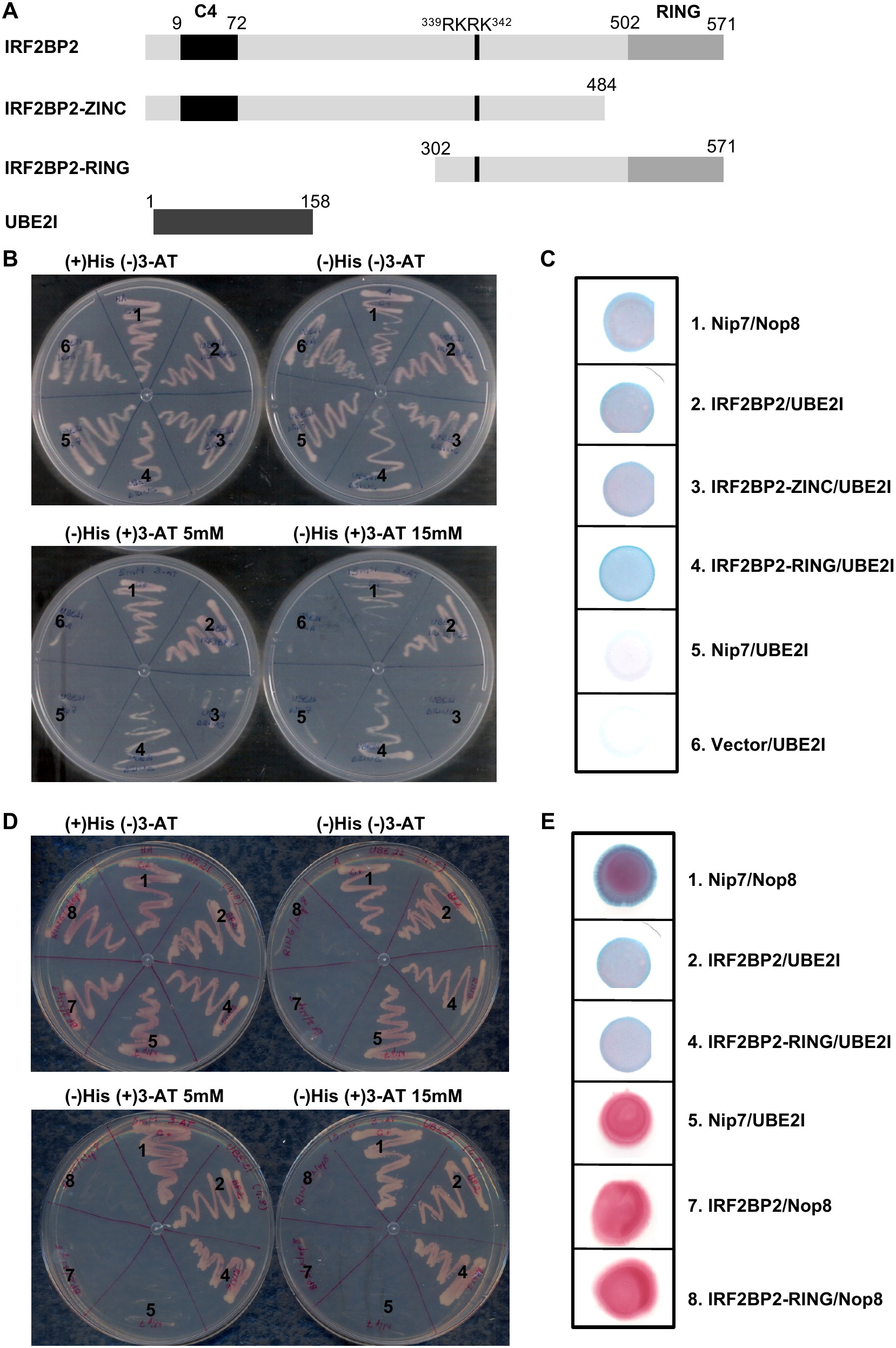
IRF2BP2 interacts with UBE2I. **(A)** Schematic representation of IRF2BP2 protein showing its conserved domains: C4 zinc finger and C3HC4 RING finger. The NLS (339RKRK342) is indicated. The two truncated proteins constructed and used in this study are represented: IRF2BP2-ZINC and IRF2BP2-RING, which are deleted for the RING and C4 zinc finger domain, respectively. The SUMO E2 ligase UBE2I is represented at the bottom. Numbers indicate the amino acids’ positions. **(B-E)**: The L40 strain, which contains the HIS3 and lacZ reporter genes, was co-transformed with vectors containing the lexA DNA binding domain fusion proteins and Gal4p activation domain fusion proteins. **(B and D)** Yeast two-hybrid interaction assay on plates containing synthetic minimal medium and 0, 5, or 15 mM 3-AT. **(C and E)** β-galactosidase filter assay. 1: positive control (pTL-Nip7p + pACT-Nop8); 2: test (pTL-IRF2BP2 + pGAD424-UBE2I); 3: test (pTL-IRF2BP2-ZINC + pGAD424-UBE2I); 4: test (pTL-IRF2BP2-RING + pGAD424-UBE2I); 5, 6, 7, and 8: negative controls (pTL-Nip7p + pGAD424-UBE2I; pTL-1 + pGAD424-UBE2I; pTL-IRF2BP2 + pACT-Nop8; pTL-IRF2BP2-RING + pACT-Nop8, respectively). All results are representative of at least three independent experiments.

The HIS3 reporter assay was performed in a medium containing high concentration of the inhibitor 3-aminotriazole (3-AT), up to 15mM. Growth under those conditions indicates a positive interaction (**Figure 1B and 1D**). The lacZ reporter gene was also evaluated, confirming the positive interaction observed (**Figure 1C and 1E**). The known interaction between Nip7p and Nop8p proteins was used as a positive control (39) (test 1 in **Figure 1B-E**). Controls represent non-interacting bait and prey proteins which were used to confirm interaction specificity (test 5 in **Figures 1B and 1C**; tests 4, 5 and 6 in **Figure 1D and 1E**). Also, the bait (IRF2BP2) with empty prey vector (not shown), and the prey (UBE2I) with empty bait vector were tested for autoactivation (test 6 in **Figure 1B and 1C**). Finaly, empty prey and bait vectors were tested to assess background activity (not shown).

Once we identified UBE2I as a possible partner of IRF2BP2, we next analyzed their localization in the cell. Immunofluorescence assays were performed using HEK293T cells transfected with UBE2I- and c-Myc-tagged IRF2BP2-expressing vectors. To simplify the nomenclature in the text, we will refer to this construct as IRF2BP2. 48 hours after transfection, cells were fixed, proteins labeled with anti-c-Myc tag and anti-UBE2I-specific antibodies, followed by Alexa488- and Rhodamine-conjugated antibodies, respectively, and observed on confocal microscopy. Our data showed that UBE2I localized in both cellular compartments, the nucleus and the cytoplasm. Whereas IRF2BP2 was mainly nuclear under those experimental conditions, as shown in the representative **Figure 2A**, but it was observed in the cytoplasm as well. Our results demonstrated that the IRB2BP2 and UBE2I proteins colocalized in the nucleus (**Figure 2A**).

**Figure 2.**
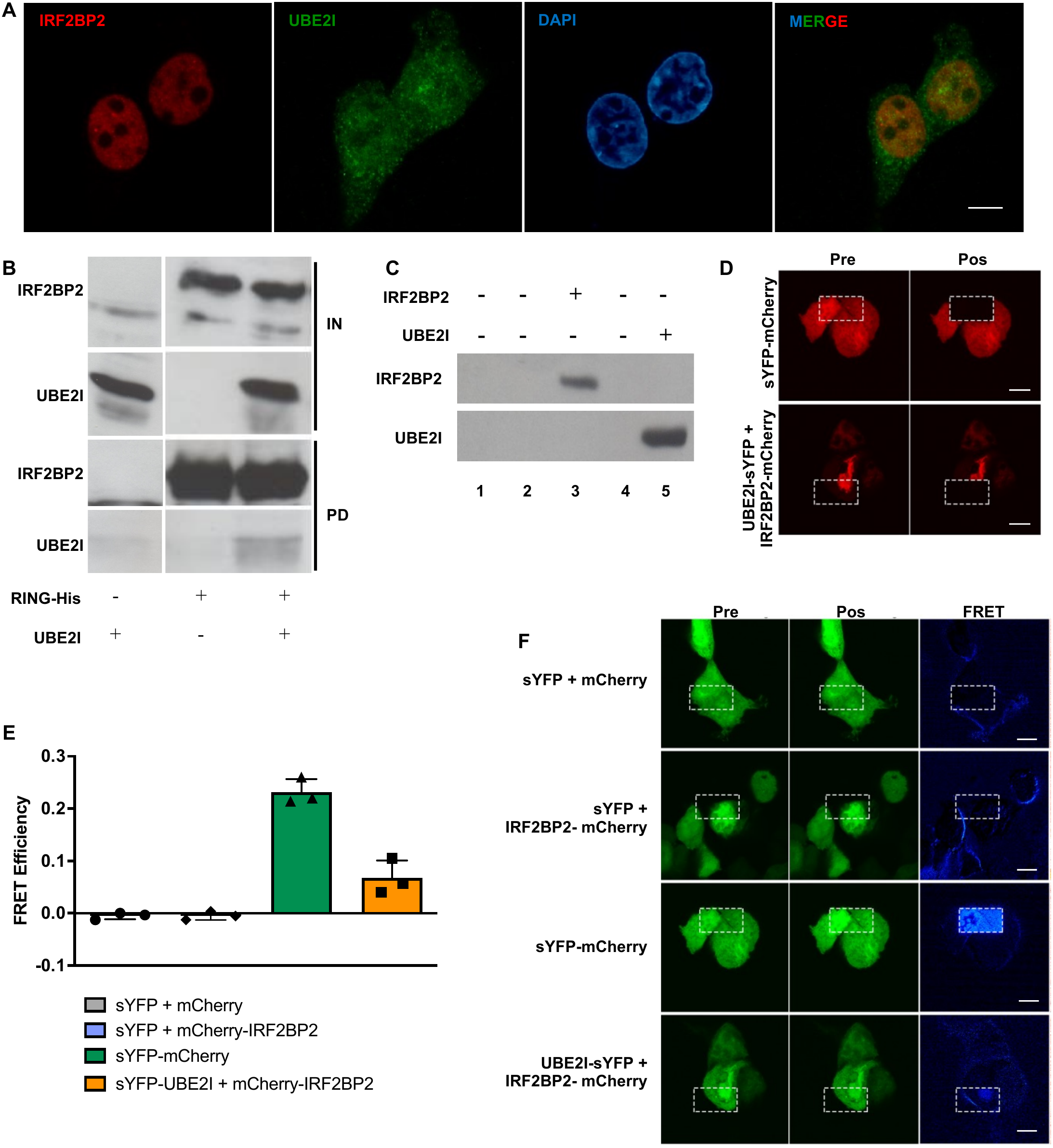
IRF2BP2 interacts with UBE2I via RING domain. **(A)** Immunofluorescence analysis of IRF2BP2 and UBE2I. HEK293T cells were co-transfected with IRF2BP2 and UBE2I and grown on glass coverslips. The cells were fixed, and the proteins were detected by specific antibodies, followed by Alexa 488- and Alexa 594-conjugated antibodies. DAPI staining indicates the nucleus position. The images were captured using confocal microscopy. **(B)** In vitro binding assay. Ni+2 beads were loaded with IRF2BP2-RING-His proteins, washed, and incubated with HEK293T extracts expressing UBE2I as indicated. The proteins were eluted from the resin by imidazole solution. Extracts and eluates were resolved by SDS-PAGE, followed by immunoblot analysis with anti-IRF2BP2 and anti-UBE2I antibodies. **(C)** HEK293T cells were transfected with codifying plasmids for IRF2BP2-mCherry or sYFP-UBE2I, as indicated in the figure. After 48 hours, cells were lysed and protein extracts obtained. Proteins were detected by specific antibodies in western blotting. 1: not transfected cells; 2: pmCherry; 3: pmCherry-IRF2BP2, 4: psYFP, 5: psYFP-UBE2I. **(D)** FRET assay by acceptor-photobleaching method. Fluorescent mCHERRY acceptor was bleached in the region of interest (ROI). **(E)** FRET efficiency pixel by pixel was calculated using three images of each condition, analyzed by FRET software. The graph represents the mean values obtained of FRET efficiency, where the value of the extracellular region (background) was subtracted. **(F)** Representative image of FRET interaction assay. mCherry-IRF2BP2 and sYFP-UBE2I proteins were produced in cotransfected HEK293T cells. After 48 hours, the cells were fixed and analyzed by confocal microscopy to verify FRET occurrence. Positive control: sYFP-mCherry. Negative controls: sYFP + mCherry; sYFP + mCherry-IRF2BP2. Test: UBE2I-sYFP + IRF2BP2-mCherry. FRET: Föster Resonance Energy Transfer; size bar = 10μm.

Next, a His pull-down assay was performed to evaluate the interaction between IRF2BP2 and UBE2I. His-tagged IRF2BP2-RING was immobilized on nickel-affinity beads and incubated with lysates from cells either expressing or not expressing recombinant UBE2I. Following extensive washing to remove unbound proteins, bound fractions were eluted and analyzed by SDS-PAGE and immunoblotting using specific antibodies. UBE2I was detected in the pull-down fraction (**Figure 2B**), suggesting an interaction with the RING domain of IRF2BP2. Control experiments using only the IRF2BP2 or only UBE2I confirmed the specificity of the binding.

To ensure the interaction between IRF2BP2 and UBE2I occurs in vivo, we used FRET (Foster Resonance Energy Transfer) assay by acceptor photobleaching method (40). This method is a powerful technique sensitive to the nanometer-range proximity (typically 1-10 nanometers) (41) and permits the determination of the occurrence of interaction between molecules. Here, we used sYFP and mCherry fluorescent proteins as donor and acceptor, respectively. To perform the assay, UBE2I-YFP and IRF2BP2-mCherry recombinant proteins were created by fusion with sYFP and mCherry, respectively. sYFP-mCherry fused protein was used as positive control. The expression of these recombinant proteins was verified by western blotting (**Figure 2C**). These recombinant proteins were produced in HEK239T cells which were then fixed and submitted for analysis by confocal microscopy to verify the FRET occurrence. Therefore, photobleaching was performed specifically on the mCherry fluorescence in a defined region of interest (ROI, **Figure 2D**). The same ROI was used in all the different analyzed combinations, always taking cytosol and nucleus included. Images of the sYFP fluorescence pre- and post-bleaching were acquired and analyzed to verify FRET occurrence. Ten images were used to quantify FRET efficiency, resulting in a score of 0.24 for sYFP-CHERRY positive control and 0.08 for UBE2I-sYFP + IRF2BP2-mCherry (**Figure 2E**). Representative images are shown in the **Figure 2F**. Cells co-expressing the single fluorescent proteins (sYFP + mCherry) represented negative controls and showed no significant increase in sYFP fluorescence (**Figure 2E and 2F**). Overall, these observations demonstrated that IRF2BP2 protein interacts with UBE2I, the SUMO-E2 ligase. Additionally, it suggests that the interaction is dependent on the RING finger of IRF2BP2 protein.

### IRF2BP2 is multiSUMOylated by SUMO1

The interaction between IRF2BP2 and UBE2I suggests that IRF2BP2 is potentially SUMOylated. Accordingly, analysis of the IRF2BP2 sequence at SUMOplot (42), found four lysine residues with high probability of being SUMO modified: lysines in the positions 566, 326, 82, and 372. To verify whether they are SUMO1 targets, all those residues were mutated to Arginine, creating IRF2BP2-KR protein which was used in the following assays.

A UBE2I dominant-negative (UBE2I-DN) was generated to perform more accurate analysis as reported by Gong (43) and Giorgino (44). The essential cysteine (Cys93) localized in the enzyme’s active site was mutated to Serine blocking its SUMO conjugation activity. Its effect on SUMOylation was observed using HEK293T cells transfected, simultaneously or not, with SUMO1-expressing plasmid. There was an intense SUMOylation when SUMO1 was overexpressed, whereas the SUMOylation was strongly decreased in the presence of UBE2I-C93S, denominated UBE2I-DN (Supporting Information **Figure S1A**).

To analyze whether IRF2BP2 is SUMOylated, an immunoprecipitation (IP) assay was performed. HEK293T cells were transfected with different combinations of plasmids expressing IRF2BP2, IRF2BP2-KR, SUMO1, and UBE2I-DN. After 48 hours, cells were harvested and lysed. IRF2BP2 was immunoprecipitated using an antibody anti-c-myc tag. After washing beads thoroughly to reduce non-specific binding, bound fractions were eluted and resolved by SDS-PAGE. The presence of SUMO1 modification was detected using an anti-SUMO1 antibody. The results showed three slower migration bands (IP: IRF2BP2, third column **Figure 3A**), suggesting that SUMO1 was added to the IRF2BP2 protein. The intensity of those three bands decreased in the presence of UBE2I-DN, highlighting that the post-translational modifications were indeed due to SUMO addition (**Figure 3A**). Besides that, no band related to SUMOylation was detected when IRF2BP2-KR was immunoprecipitated in the presence of SUMO1 (sixth column, **Figure 3B**), suggesting that the lysine residues targeted for SUMOylation were mutated. Since it is known SUMO1 does not form poliSUMO chains (45, 46), these data indicate that SUMO1 can modify IRF2BP2 protein at three Lysine residues. Our data indicates that IRF2BP2 is multiSUMOylated.

**Figure 3.**
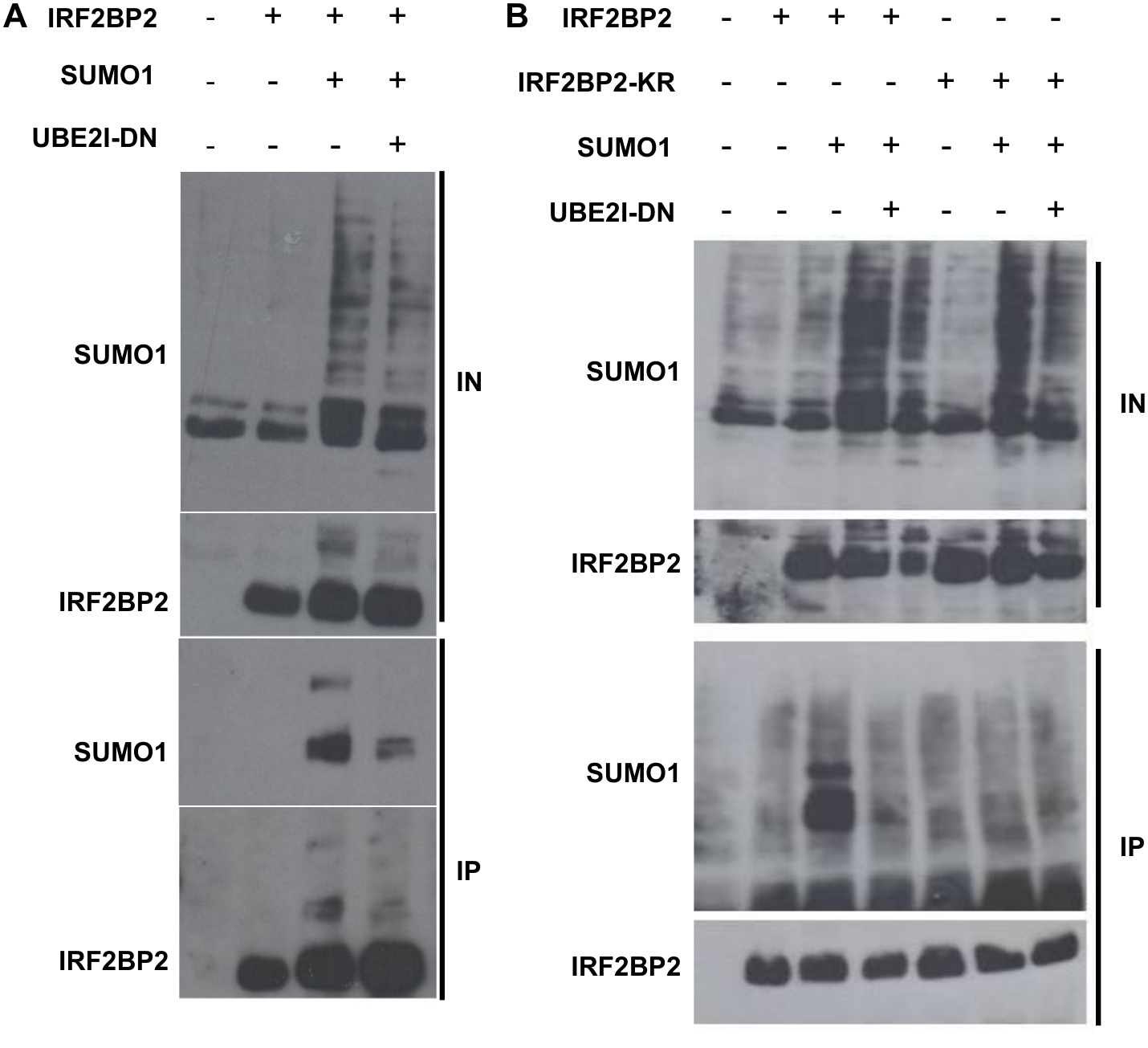
IRF2BP2 is multiSUMOylated by SUMO1. HEK293T cells were transfected with expressing plasmids as indicated. After 48 hours, protein extracts were obtained and quantified. Equal quantities of total protein were loaded and IRF2BP2 was immunoprecipitated. Inputs and elutions were analyzed by SDS-PAGE and proteins detected by anti-SUMO1 and anti-IRF2BP2 antibodies. **(A)** Cells transfected with IRF2BP2, SUMO1, and UBE2I-DN. **(B)** Cells transfected with IRF2BP2, IRF2BP2-KR, SUMO1 and UBE2I-DN. In both, cells transfected with empty vector were used as negative controls. Results are representative of at least three independent experiments.

### IRF2BP2 protein is predominantly nuclear in the S phase of the cell cycle

Once our data showed that IRF2BP2 is SUMO1 modified, we next investigated the impact of this modification in its function. It has been well reported in the literature that SUMO1 conjugation is dynamic, reversible, and modulates several cellular processes, including protein stability, subcellular localization, and protein-protein interactions (46, 47).

Since we observed IRF2BP2 in the nucleus and cytoplasm, we decided to investigate IRF2BP2 localization during the cell cycle to further check whether SUMO1 has any effect on its localization. HEK293T cells were transfected with IRF2BP2 and, after 30 hours, submitted to thymidine synchronization for 18 hours. Following this time, denominated time zero (T0), the cell cycle was released. The cells were harvested at different time points, stained with propidium iodide, and subsequently analyzed by flow cytometry. The histograms showed that the thymidine synchronization arrested 90% of the cells at the G1/S boundary (**Figure 4A**). Simultaneously, cells were fixed, and IRF2BP2 was labeled with anti-c-myc tag antibody followed by Rhodamine-conjugated antibody. The nucleus was identified by DAPI. The cells were observed on microscopy and images obtained. They showed that IRF2BP2 localization changed throughout the cell cycle. Interestingly, during the S phase (0 and 3 hours, **Figure 4B**), IRF2BP2 protein is mainly nuclear, while in the other phases, it was observed in both the cytoplasm and the nucleus (6, 12, and 24 hours; **Figure 4B**). These results demonstrate that IRF2BP2 subcellular localization is regulated during the cell cycle. Its predominant presence in the nucleus at the S phase suggests an important role at this stage. However, our primary focus was to investigate whether SUMO1 contributes to determining the nuclear localization of IRF2BP2.

**Figure 4.**
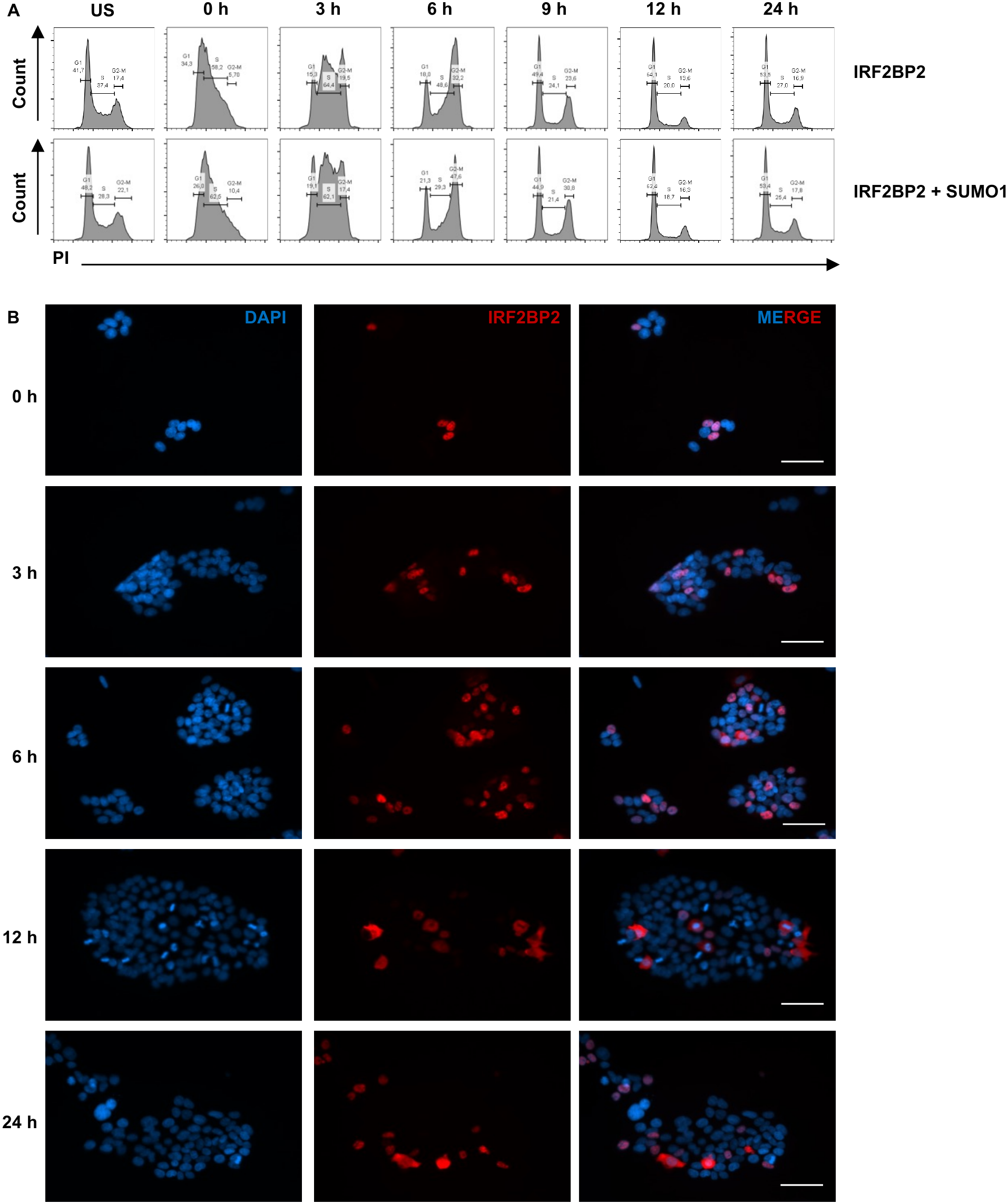
IRF2BP2 protein is predominantly nuclear in the S phase of the cell cycle. **(A-B)** HEK293T cells co-transfected with IRF2BP2 and SUMO1 or only IRF2BP2, as indicated, were submitted to thymidine synchronization (2.5 mM) for 18 hours. After this time, the cells were washed, and the cell cycle was released (= Time 0 hours). Unsynchronized and synchronized cells at different time points were fixed. **(A)** Cells were stained with Propidium iodide and analyzed by flow cytometry. Cell population frequencies in the G1, S, and G2/M phases were shown in each histogram across the time (hours). **(B)** Cells obtained at different time points (hours) were fixed, and IRF2BP2 was identified using anti-c-Myc tag antibody followed by Alexa 594-conjugated antibody. The cells were observed at Zeiss microscope. Results are representative of at least three independent experiments. US: Unsynchronized cells. Size bar = 20μm.

### SUMO1 enhances IRF2BP2 nuclear localization

The IRF2BP2 protein has been described as a nuclear protein (1, 5, 6, 48). A nuclear localization signal (NLS) was identified (6), along with a phosphorylation site near the NLS, which is also important for cytoplasm-nucleus translocation (48). However, IRF2BP2 localization has also been reported in both compartments, cytoplasm and nucleus, of myoblasts (48). Consistent with this, our current data revealed IRF2BP2 in both compartments and suggest that it can translocate between the nucleus and cytoplasm.

To further compare the localization of IRF2BP2 and IRF2BP2-KR, we performed immunofluorescence using HEK293T cells transfected with the respective constructs. Cells were fixed, stained with anti-c-myc followed by a rhodamine-conjugated secondary antibody. The nucleus was visualized with DAPI. Interestingly, while IRF2BP2 displays a homogeneous nuclear distribution, IRF2BP2-KR accumulated in distinct nuclear foci. Quantification of fifty cells from each condition confirmed this observation. IRF2BP2 showed a dispersed nuclear pattern, while IRF2BP2-KR adopted a punctate nuclear distribution (**Figure 5A and B**), suggesting that SUMO1 modulates the IRF2BP2 nuclear distribution which might impact its interaction with partners and, consequently, its function.

**Figure 5.**
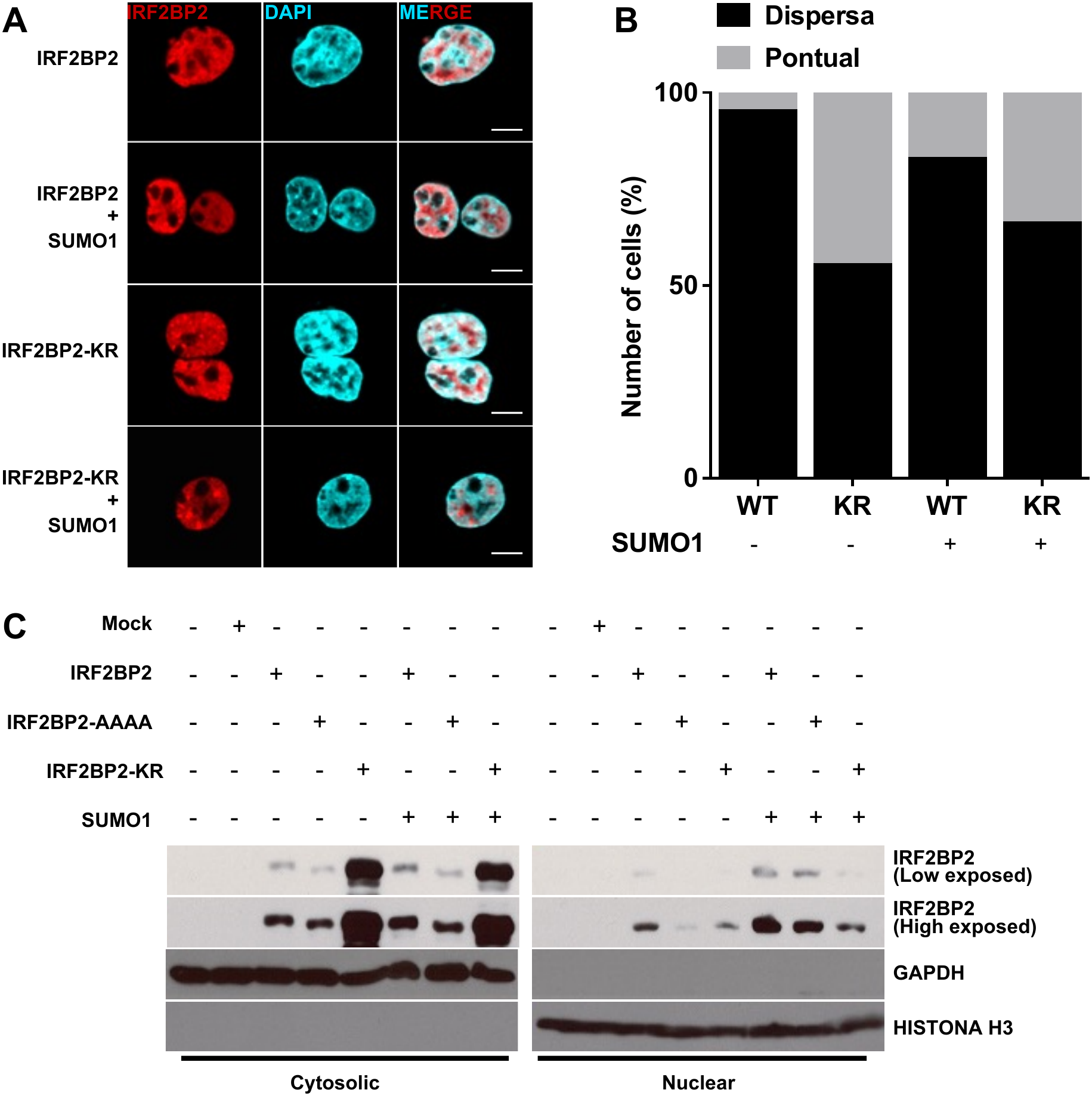
SUMO1 enhances IRF2BP2 nuclear localization. **(A-B) (A)** HEK293T cells were transfected with plasmids codifying IRF2BP2, IRF2BP2-KR, and SUMO1, simultaneously or not, and fixed after 48 hours. Immunostaining identified IRF2BP2 by anti-c-Myc tag followed by Alexa 594-conjugated antibody. DAPI indicates nucleus position. Images were acquired at Zeiss microscope. **(B)** Fifty cells of each condition were blindly counted, considering the type of IRF2BP2 nuclear distribution as dispersed or punctual. **(C)** HEK293T cells were transfected with codifying plasmids for IRF2BP2, IRF2BP2-KR, or IRF2BP2-AAAA, in the presence or absence of SUMO1, as indicated in the figure. After 48 hours, the cells were lysed, and cytosolic and nuclear extracts were obtained. Proteins were resolved by SDS-PAGE and analyzed by immunoblotting with an anti-c-myc antibody. Mock cells transfected with the empty vector served as negative controls. Purity of the extracts was assessed using GAPDH and Histone H3, which are specific markers of cytoplasm and nucleus, respectively. Mock: HEK293T transfected with empty vectors. Size bar = 10μm.

To investigate this hypothesis, we used IRF2BP2-AAAA, which carries a mutated NLS and is predominantly cytoplasmic (6), to analyze protein localization in the context of SUMOylation. IRF2BP2, IRF2BP2-KR, or IRF2BP2-AAAA were expressed in HEK293T cells with or without SUMO1. Forty-eight hours after transfection, cells were harvested and lysed to obtain cytosolic and nuclear fractions. Equal amounts of total protein were resolved by SDS-PAGE. Nuclear and cytosolic fraction purity was confirmed using anti-GAPDH (cytoplasmic marker) and anti-Histone H3 (nuclear marker). Anti-c-Myc tag antibody was used to detect the IRF2BP2 constructs. We observed IRF2BP2 in both nuclear and cytoplasmic fractions. Notably, nuclear levels of IRF2BP2 increased in the presence of SUMO1 (**Figure 5C**). In contrast, the IRF2BP2-AAAA protein was highly expressed but restricted to the cytoplasm, consistent with previous findings (6). Only trace amounts were detected in the nucleus, likely reflecting minor cytoplasmic-to-nuclear leakage of the protein (**Figure 5C**). Interestingly, the non-SUMOylated mutant protein IRF2BP2-KR was present in both compartments, suggesting that, beyond SUMOylation, additional mechanisms contribute to IRF2BP2 nuclear translocation. Taken together, these results demonstrate that SUMO1 directly modulates the availability and spatial organization of IRF2BP2 within the nucleus. By shifting the protein from a restricted, punctate accumulation (foci) to a more homogeneous, dispersed distribution, and by increasing its total nuclear level, SUMOylation likely regulates the dynamic interaction of IRF2BP2 with specific nuclear partners, thereby modulating its transcriptional regulatory functions.

### Importin β1-dependent nuclear import and euchromatin localization of IRF2BP2

IRF2BP2 functions have been so far primarily described within the nucleus(1, 6, 10). Consistent with that, our data suggest that SUMO1 conjugation contributes to regulate IRF2BP2 subcellular localization. It well demonstrated that SUMO1 can influence nucleocytoplasmic trafficking and play a critical role in NLS-dependent nuclear import. In addition, SUMO1 conjugation enhances nuclear accumulation by masking nuclear export signals (NES), supporting nuclear retention, or altering protein interactions, rather than directly affecting nuclear import kinetics (49). The nuclear transport of macromolecules is primarily mediated by the karyopherin (Importin)-β family which depend on the presence of NLS or NES to determine subcellular distribution. Importins recognize and bind to NLS-containing proteins, facilitating their transport into the nucleus through the nuclear pore complex. Classical NLS motifs are typically recognized by the importin-α/β heterodimer (50).

To investigate the potential mechanism of IRF2BP2 nuclear import via importin pathways, we employed the inhibitors Ivermectin and INI-43 to assess IRF2BP2 subcellular localization. Ivermectin inhibits importin α/β1-dependent nuclear transport (51), while INI-43 specifically targets importin β1 (52). HEK293T cells transfected with constructs overexpressing IRF2BP2 and SUMO1 were used in the assay. Forty-eight hours post-transfection, cells were treated with the inhibitors (10 µM) for 1 hour and fixed to perform immunofluorescence. IRF2BP2 was detected using anti-c-Myc tag antibody, followed by Alexa 594-conjugated secondary antibody. Nucleus position was determined by using DAPI, and cellular localization was examined by fluorescence microscopy. Treatment with INI-43 led to complete cytoplasmic retention of IRF2BP2, particularly in those SUMO1-overexpressing cells, whereas ivermectin treatment resulted in both nuclear and cytoplasmic localization. This differential effect indicates that IRF2BP2 nuclear import depends on the importin β-mediated pathway, which is effectively inhibited by INI-43. In contrast, ivermectin, which preferentially targets the importin α/β1 complex, only partially interferes with IRF2BP2 translocation, suggesting that importin α is not essential for this process. Together, these results support a model in which SUMO1 modification promotes importin β1-dependent nuclear import of IRF2BP2, independently of the classical import α/β1 pathway. Interestingly, SUMO1 is also able to revert ivermectin effect, but not the INI43 effect (**Figure 6A**).

**Figure 6.**
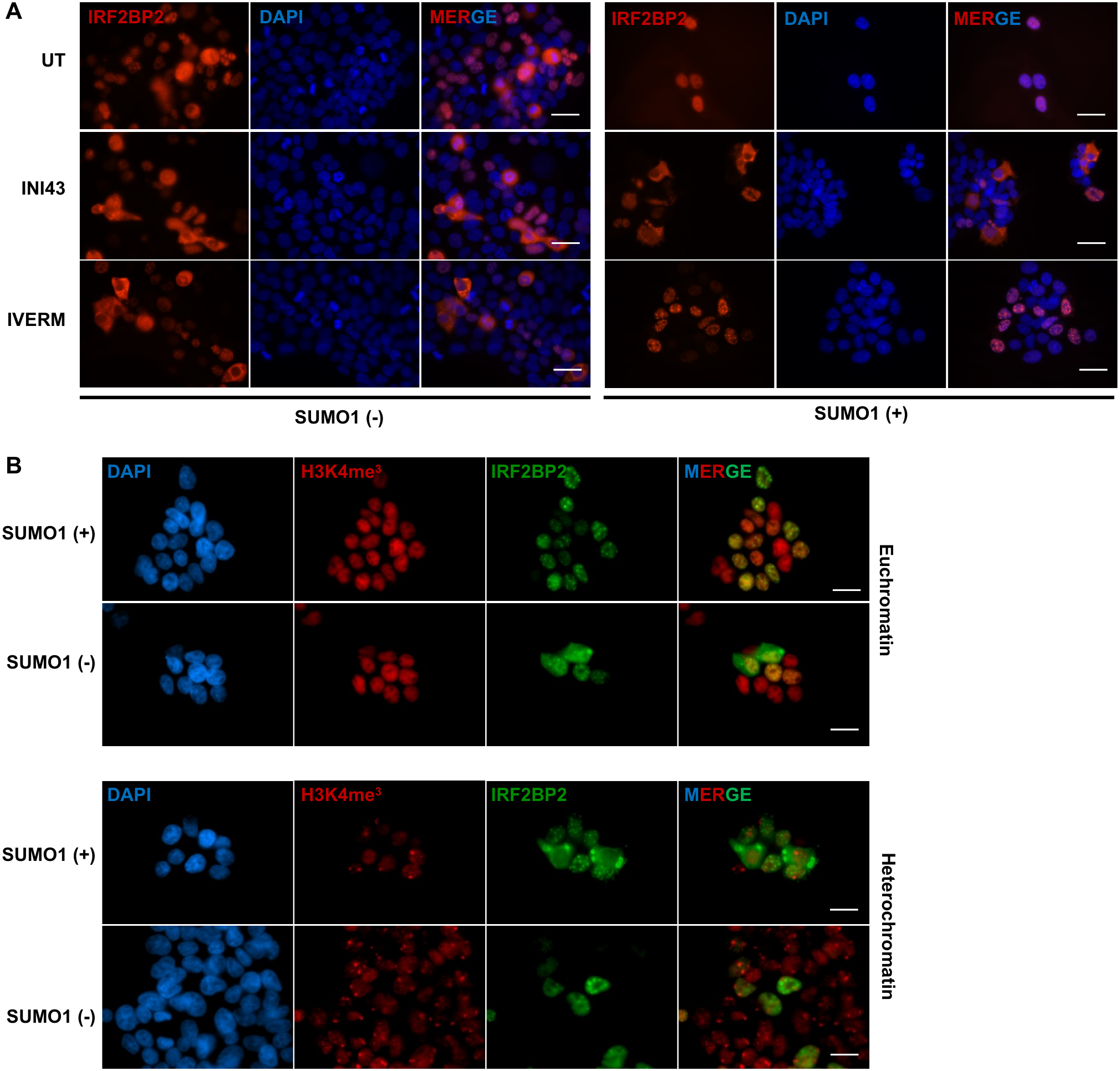
Importin β1-dependent nuclear import and euchromatin localization of IRF2BP2. **(A)** HEK293T cells expressing IRF2BP2, with or without SUMO1, were treated, 48 hours after transfection, with INI43 or Ivermectin (both at 10 µM) for 1 hour. Cells were fixed, and IRF2BP2 was labeled with anti-c-myc tag followed by Alexa 594-conjugated antibody. Untreated (UT) cells were used as negative control. **(B)** HEK293T cells expressing IRF2BP2, with or without SUMO1, were fixed, and the proteins were detected with specific antibodies. IRF2BP2 was identified using anti-c-myc tag antibody, followed by Alexa 488-conjugated-antibody. Trimethylated-Lysine 4 Histone H3 and Trimethylated-Lysine 27 Histone H3 were identified by specific antibodies to detect the respective modifications, followed by Alexa 594-conjugated antibody. DAPI indicates nucleus position. Images were acquired at Zeiss microscope. Size bar = 20μm.

Lastly, we examined the intranuclear localization of IRF2BP2. Cells were co-transfected with vectors overexpressing IRF2BP2 and SUMO1. After 48 hours, cells were fixed, and IRF2BP2 was detected using anti-c-myc tag antibody followed by an Alexa-488-conjugated secondary antibody. Immunofluorescence staining was simultaneously performed for the euchromatin marker H3K4me3 and the heterochromatin marker H3K27me3, followed by Alexa 594-conjugated antibody. Images acquired revealed that IRF2BP2 predominantly localized to the euchromatin region within the nucleus (**Figure 6B**, upper panel). Notably, IRF2BP2 did not colocalize with heterochromatin regions (**Figure 6B**, bottom panel). These results suggest that IRF2BP2 nuclear import is importin β-dependent and, once in the nucleus, IRF2BP2 preferentially localizes to transcriptionally active euchromatin regions, consistent with its proposed regulatory roles.

### SUMOylation enhances IRF2BP2 protein stability

SUMO1 and ubiquitin (Ub) are covalently attached to lysine residues on numerous proteins. SUMO-acceptor lysine residues are often shared with ubiquitin. However, these modifications can exert opposing regulatory effects (53). Because SUMOylation and ubiquitination on the same residue are mutually exclusive, competition between them can influence protein-protein interactions and/or protein stability (54). In most mammalian cells, the ubiquitin-proteasome pathway is responsible for most of the intracellular protein degradation (55). Therefore, we next addressed whether SUMO1 modulates IRF2BP2 stability, and consequently its functional lifespan.

First, we assessed IRF2BP2 protein levels in the presence of the classical proteasome inhibitor MG132. HEK293T cells were transfected with plasmids encoding IRF2BP2 or the SUMO-deficient mutant IRF2BP2-KR, as indicated. After 48 hours, cells were treated with MG132 (20 µM) for 24 hours. No accumulation of IRF2BP2 was observed following MG132 treatment (left, **Figure 7A**), indicating that IRF2BP2 is not degraded by proteasome. Similar results were obtained for the SUMO-deficient protein (right, **Figure 7A**), suggesting that the absence of SUMO1 does not affect proteasome-mediated degradation of IRF2BP2.

**Figure 7.**
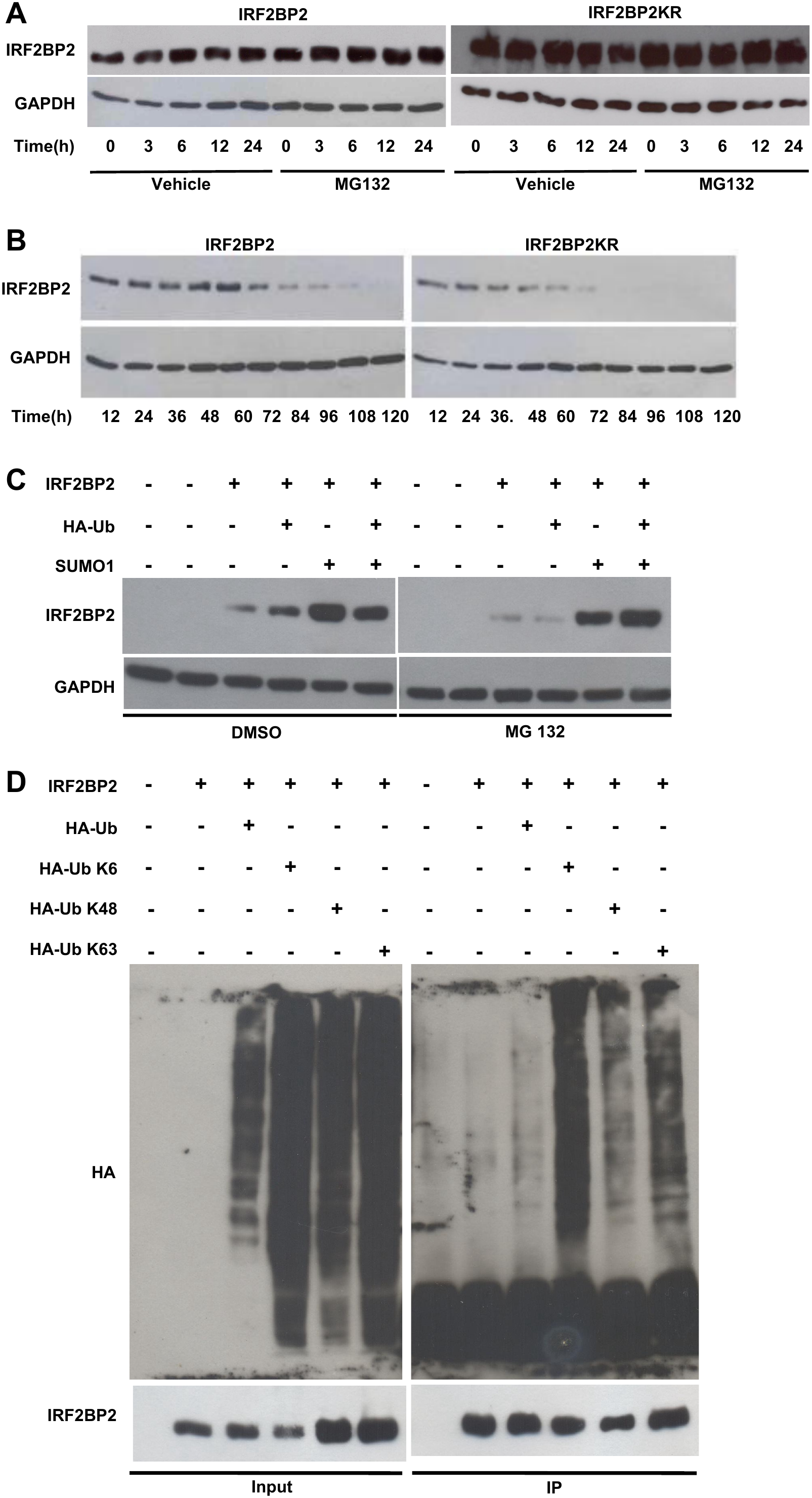
SUMOylation enhances IRF2BP2 protein stability. **(A-B)** HEK293T cells were transfected with codifying plasmids for IRF2BP2 or IRF2BP2-KR, as indicated. After 48 hours, cells were treated with MG132 proteasome inhibitor (20 µM). Protein extracts were obtained at different time points (hours) during the treatment. **(A)** MG132 treatment performed for 24 hours. **(B)** MG132 treatment performed for 120 hours. Protein extracts were obtained every 12 hours. **(C)** HEK293T cells were transfected with combinations of codifying plasmids for IRF2BP2, SUMO1, and HA-tagged Ubiquitin (Ub), as indicated. After 48 hours, cells were treated with MG132 (20 µM) for 2 hours. Protein extracts were obtained for each condition, resolved by SDS-PAGE. **(D)** HEK293T cells were transfected with codifying plasmids for IRF2BP2, HA-Ub, and the Ubiquitin variants HA-Ub K6, HA-Ub K48, HA-Ub K63, as indicated in the figure. After 48 hours, protein extracts were obtained. IRF2BP2 was immunoprecipitated using anti-c-myc-tag. Inputs and elutions were analyzed by western blot. Antibodies used to detect respective proteins: Anti-IRF2BP2, anti-HA, and anti-GAPDH antibodies.

Next, we examined IRF2BP2 protein levels over a 120-hour period. Protein extracts from HEK293T cells expressing either IRF2BP2 or the SUMO-deficient IRF2BP2-KR proteins were analyzed by western blotting. Interestingly, IRF2BP2-KR levels declined earlier than those of wild-type IRF2BP2 (**Figure 7B**), suggesting that SUMOylation may enhance IRF2BP2 stability. Corroborating this finding, the presence of SUMO1 increased IRF2BP2 protein levels independently of ubiquitin (**Figure 7C**). IRF2BP2 was co-expressed with HA-tagged ubiquitin or SUMO1 in HEK293T cells, as indicated and equal amounts of total protein analyzed. Consistent with our previous observations (**Figure 7A and 7B**), MG132 treatment did not alter IRF2BP2 abundance (**Figure 7C**).

Since our results showed IRF2BP2 undergoes ubiquitination, but it does not determine its proteasome degradation, we next investigated what kind of ubiquitin chains are conjugated to IRF2BP2. The functional outcome of ubiquitination depends on the type of chain linkage (56). To address this, HA-tagged ubiquitin was co-expressed in HEK293T cells, and IRF2BP2 was immunoprecipitated using an anti–c-Myc antibody. Western blotting with anti-HA revealed ubiquitin conjugation to IRF2BP2, confirming the formation of ubiquitin chains (**Figure 7D**). Followed that, we investigated which ubiquitin chain is preferentially formed on IRF2BP2. Ubiquitin contains seven lysine residues (K6, K11, K27, K29, K33, K48, K63), enabling the formation of distinct linkages. K48-linked chains are the most abundant in cells and typically target substrates for proteasomal degradation (Sun and Zhang, 2022). K63-linked chains have non-proteolytic roles, including regulating intracellular trafficking and DNA repair. Other linkages are less common; among them, K6-linked chains remain poorly understood but have been implicated in DNA damage responses and mitochondrial quality control (57).

To determine the type of ubiquitin chain formed on IRF2BP2, we used single-lysine ubiquitin mutants, each capable of forming only one linkage type. Equal amounts of protein were used for IRF2BP2 immunoprecipitation. Comparison of K6-, K48-, and K63-linked chains showed that IRF2BP2 is predominantly modified by K6-linked ubiquitin, which is considered non-proteolytic (**Figure 7D**). Supporting our conclusion that IRF2BP2 is not degraded via proteasomal pathway, low levels of K48-linked chains were detected (**Figure 7D**). The biological significance of K6-linked ubiquitination on IRF2BP2 remains to be elucidated and represents an important direction for future studies. Moreover, which alternative pathway mediates IRF2BP2 degradation remains unknown and warrants further investigation.

## DISCUSSION

IRF2BP2 is a multifunctional protein that primarily acts as a transcriptional coregulator, functioning as either a repressor or an activator depending on cellular context (7, 11). Although IRF2BP2 regulates the expression of genes involved in diverse biological processes, the mechanisms controlling its own regulation remain unclear. To investigate these mechanisms, we performed a yeast two-hybrid screen using full-length IRF2BP2 as bait and identified UBE2I as a potential interacting partner. UBE2I is the sole known E2 SUMO-conjugating enzyme and plays a central role in substrate recognition and SUMO transfer during protein SUMOylation (45, 58). We subsequently showed that they colocalize and validated the interaction through pull-down, immunoprecipitation, and FRET assays. We also demonstrated this interaction depends on the RING-domain of IRF2BP2. Importantly, SUMOylation assays revealed three distinct slower-migrating IRF2BP2 protein bands consistent with SUMO1 conjugation, which were markedly reduced or nearly abolished upon expression of a dominant-negative UBE2I mutant (UBE2I-DN). Together these findings identify UBE2I as an IRF2BP2-interacting protein and support its involvement in the SUMO-dependent post-translational regulation of IRF2BP2.

To further characterize IRF2BP2 SUMOylation, we identified four high-probability SUMOylation sites using SUMOsp (42): K82, K326, K372, and K566. Substitution of these lysine residues with arginine (our IRF2BP2-KR) markedly reduced SUMO1 conjugation, supporting their contribution to IRF2BP2 SUMOylation. Previous studies a conserved C-terminal lysine as SUMOylation site correspondent to K566 in zebrafish Irf2bp2 (5) and in human IRF2BP2 (38). Our findings extend these observations by suggesting that additional lysine residues contribute to IRF2BP2 SUMOylation, highlighting a potentially more complex pattern of post-translational regulation than previously identified.

The dynamic balance between SUMOylated and unSUMOylated states is essential for regulating diverse cellular processes. Its dysregulation has been implicated in several pathological conditions, including cancer, cardiovascular disorders, and autoimmune diseases (31, 59-61). Among its regulatory functions, SUMOylation can influence protein subcellular localization (47, 62). In this context, we observed that IRF2BP2 exhibits cell cycle-dependent subcellular distribution, with predominant nuclear localization during S phase and detectable cytoplasmatic localization at others phases as well. Although the functional significance of this redistribution remains to be clarified, these observations suggest that IRF2BP2 localization is dynamically regulated throughout the cell cycle.

We therefore investigated whether SUMOylation contributes to IRF2BP2 nuclear localization. Subcellular fractioning revealed increased nuclear accumulation of IRF2BP2 upon SUMO1 overexpression., supporting a role for SUMOylation in promoting its nuclear localization. Interestingly, the SUMO-deficient IRF2BP2-KR mutant retained nuclear localization but exhibited a distinct punctate distribution, in contrast to the more diffuse nuclear pattern observed for wild-type IRF2BP2. These observations raise the possibility that SUMOylation contributes to the IRF2BP2 intranuclear organization, in addition to promoting its nuclear accumulation. Previous work from our group demonstrated that mutation of the nuclear localization signal (NLS) restricts IRF2BP2-AAAA to the cytoplasm (6), highlighting the importance of an intact NLS for the nuclear import. Moreover, UBE2I has been reported in cytoplasm, nuclear membrane, and nucleoplasm across different cell types (63-65), consistent with the possibility that IRF2BP2 SUMOylation may occur in multiple subcellular compartments. Collectively, these findings suggest that SUMOylation promotes IRF2BP2 nuclear accumulation, whereas its NLS remains a critical determinant of nuclear localization.

To further investigate the mechanism underlying IRF2BP2 nuclear localization, we examined the involvement of importin-mediated transport. Importins facilitate the nuclear import of numerous proteins, including those harboring classical NLSs. We therefore employed two pharmacological inhibitors, ivermectin and INI43, which interfere with importin α/β1-dependent and importin β1-dependent nuclear transport, respectively(51, 52). Immunofluorescence analysis revealed that INI43 treatment promoted cytoplasmatic retention of IRF2BP2, an effect that was particularly pronounced upon SUMO1 overexpression. In contrast, ivermectin treatment did not produce a comparable redistribution, with IRF2BP2 remaining detectable in both nuclear and cytoplasmatic compartments. Consistent with our previous observations, SUMO1 overexpression favored nuclear accumulation of IRF2BP2 in untreated cells. Collectively, these findings support the involvement of importin β1-dependent transport in IRF2BP2 nuclear localization and suggest that SUMOylation facilitates this process in cooperation with its NLS.

Beyond nuclear import, we investigated the intranuclear distribution of IRF2BP2 by analyzing its association with chromatin regions marked by H3K4me3 and H3K27me3, which are typically associated with transcriptionally active and Polycomb-repressed chromatin, respectively. IRF2BP2 preferentially colocalized with H3K4me3-enriched nuclear regions, an association that was emphasized upon SUMO1 overexpression. Given the established role of IRF2BP2 as a transcriptional coregulator, these observations suggest that SUMOylation may influence its spatial association with transcriptionally active chromatin. Such regulation could have implications for the diverse biological processes in which IRF2BP2 participates, including cell differentiation, immune responses, inflammation, and tumorigenesis. However, whether SUMOylation directly modulates IRF2BP2 chromatin recruitment or its transcriptional regulatory activity remains to be determined.

We further demonstrate that SUMOylation enhances IRF2BP2 protein stability, contributing to its increased abundance. In parallel, we identified ubiquitination of IRF2BP2, predominantly involving K6-linked ubiquitin chains. Although the functional significance of this modification remains to be elucidated, our findings suggest that IRF2BP2 ubiquitination may participate in regulatory processes beyond canonical proteasomal degradation. Taken together, our findings reveal SUMOylation as an important regulatory mechanism influencing IRF2BP2 stability, nuclear trafficking, and intranuclear distribution. This work expands our understanding of the post-translational regulation of IRF2BP2 and provides a foundation for investigating how these modifications influence its context-dependent functions as a transcriptional coregulator.

## MATERIAL AND METHODS

### Cell culture and transfection

HEK293T cells were maintained in Dulbecco’s modified medium supplemented with 10% fetal bovine serum (FBS), L-glutamine, penicillin-streptomycin, essential and nonessential amino acids, sodium pyruvate, vitamins, HEPES and β-mercaptoethanol (All from Invitrogen) at 37°C and 5% CO_2_ in a humidified environment. HEK293T cells (1,5 x 10^7^) were transfected by the calcium phosphate method (CaCl2: 250 mM; HBS Buffer: 140 mM NaCl, 5 mM KCl, 750 µM Na2HPO4, 6 mM dextrose, 25 mM HEPES) with 10 µg DNA of distinctive plasmids as described for the different experiments. Forty-eight hours after transfection, cells were collected and lysed with specific buffers depending on the assay.

### Plasmid construction

In this study, it was used the human IRF2BP2 isoform B, which was c-myc-His tagged and constructed as described by Carneiro (6). UBE2I and SUMO1 were each one inserted into pcDNA5 vector. The Lysine residues 566, 326, 82, and 372 were predicted with high probability of being SUMO modified by GPS-SUMO (RRID:SCR_018261). They were mutated to Arginine (by Genescript), creating IRF2BP2-KR which was inserted into the pcDNA4 vector. The negative dominant pcDNA5-UBE2I-C93S was constructed by site-directed mutagenesis to mutate Cysteine in position 93 (TGC) to Serine (TCC) using the primers described in Table S2 and GENEART Site-Directed Mutagenesis System (Invitrogen) as recommended by the manufacturer. To perform FRET assays, IRF2BP2 was cloned into pmCherry-N1 and UBE2I, into psYFP-N1. Both plasmids pmCherry-N1 and psYFP-N1 were kindly donated by Andres Cardozo Gizzi and pYFP-Cherry was acquired from Clontech. All plasmids used in this work and cloning details are listed in **Table S1**, and all primers used to construct them in **Table S2**. pHA-Ub, pHA-Ub-K6, pHA-Ub-K48 and pHA-Ub-K63 were kindly gifted by Dr. Marcelo Alex de Carvalho (IFRJ, Brazil).

### Antibodies and reagents

All Antibodies and reagents are listed in **Table S3**.

### Yeast two-hybrid assay

A large-scale screening was performed using IRF2BP2 as bait. The L40 strain (Hollenberg, 1995) containing the pTL-IRF2BP2 plasmid was transformed with an Epstein-Barr virus (EBV)-transformed human B cell cDNA library fused to the activation domain of Gal4p in the pACT vector (Clontech). Positive clones were selected on plates containing YNB (Yeast nitrogen base) medium supplemented with adenine and 3 mM 3-amino-triazole (3-AT), followed by β-galactosidase assays (66). Plasmid DNA was extracted from the positive clones, transformed into E. coli XL-1 Blue (Stratagene) for amplification and subjected to DNA sequencing analysis (67). For further interaction analyses, the L40 strain was cotransformed with pTL-IRF2BP2/pACT-UBE2I, pTL-IRF2BP2-ZINC/pACT-UBE2I, pTL-IRF2BP2-RING/pACT-UBE2I. pTL-Nip7/pACT-Nop8(39) was used as a positive control, pTL-Nip7/pACT-UBE2I and pTL-1/pACT-UBE2I were used as negative controls. The test and control strains were grown in YNB medium containing either histidine or 5 and 15 mM 3-AT. Filter assays to test β-galactosidase expression were also performed.

### His-Pull-Down Assay

HEK293T cells (1,5 x 10^7^) were transfected with pcDNA4-IRF2BP2-RING, which contains c-myc and 6xHis (His) tags, or with pcDNA5-UBE2I. After 48 hours, protein extracts were obtained using lysis buffers: 20 mM Tris-Cl pH=7,4, 150 mM NaCl, 0,5% Nonidet P-40, 1 mM EDTA, 500 µM PMSF, protease inhibitor cocktail. The second buffer was modified with 2% Triton X-100 and 0.5% SDS. Then, Ni+2-NTA resin (Thermo Fisher Scientific) was incubated with the extract containing IRF2BP2-RING for 2 hours at 4°C. Subsequently, the beads loaded with the proteins were washed with the first lysis buffer and incubated for an additional 2 hours at 4°C with the extract containing UBE2I or not. The beads were washed, and the proteins were eluted in a buffer containing imidazole (300 mM). Samples of extracts and eluted were resolved by SDS-PAGE followed by immunoblot with anti-c-Myc tag and anti-UBE2I antibodies.

### FRET

The fluorescent proteins sYFP and mCherry were used as donor-acceptor pair to perform Foster Resonance Energy Transfer (FRET). HEK293T cells were cotransfected with UBE2I-YFP and IRF2BP2-CHERRY. After 48 hours, the cells were fixed and subjected to FRET analysis. An area of ROI was delimited (dotted line), including cytoplasm and nucleus, and the fluorescence of the acceptor protein in this area was photobleached. The donor protein’s fluorescence was observed, and the images were acquired before and after the photobleaching by confocal microscopy. These images were analyzed to verify FRET occurrence between mCherry and sYFP. The FRET efficiency was obtained by FV10-ASW 2.1 software. To quantify FRET, 4 images from each condition were analyzed. The background value was subtracted from the medium value of FRET and then plotted in a graph. HEK293T cells transfected with sYFP-mCherry fusion protein were used as positive control. HEK293T cells transfected with sYFP + mCherry or sYFP + IRF2BP2-mCherry proteins were used as negative controls.

### Immunoprecipitation assay

HEK293T cells (1,5 x 10^7^), maintained on dishes (10 cm), were cotransfected by the calcium phosphate method with pcDNA4-IRF2BP2 or pcDNA4-IRF2BP2-KR (pcDNA4-IRF2BP2-K82-326-372-566R) and pcDNA5-SUMO1 and, or pcDNA5-UBE2I-DN, as indicated in the different assays. Forty-eight hours later, total protein extracts were obtained using a cell scraper in lysis buffer containing protease inhibitors, followed by gentle sonication. All buffers, lysis and wash, used in IP assays were described by Hara and collegues (68). Immunoprecipitation was performed using Protein A-Sepharose 4B (GE Healthcare) using 2μg antibody anti-c-Myc-tag.

### Nuclear/Cytosolic fractionation

The nuclear and cytosolic fractions were obtained from HEK293T cells transfected with pcDNA4-IRF2BP2, pcDNA4-IRF2BP2-KR (pcDNA4-IRF2BP2-K82-326-372-566R), pcDNA4-IRF2BP2-AAAA (MutNLS), and pcDNA5-SUMO1, as indicated in the figures. The cells (2 x 10^7^) were harvested and incubated with 1 mL of hypotonic buffer (10 mM Tris, pH7.5; 10 mM KCl, 0.1 mM EDTA, 1.5 mM MgCl2, 10 mM iodoacetamide) for 30 minutes on ice. The nuclear fraction was recovered by centrifugation and resuspended in 1 mL of hypertonic buffer (20 mM Tris, pH8.0; 0.4 M NaCl, 0.1 mM EDTA, 1.5 mM MgCl2, 10 mM iodoacetamide, 25% glycerol), followed by incubation on ice for 30 minutes.

### Western blotting

HEK293T protein extracts, inputs, and eluted proteins from the different assays performed were incubated at 100°C for 10 minutes. Total protein from cells was obtained in cell lysis buffer (40 mM Tris pH 7.5; 60 mM NaPPi, 10 mM EDTA, 5% SDS) and incubated at 100°C for 10 minutes. The protein samples were loaded and separated by SDS-PAGE. After that, the proteins were transferred onto a nitrocellulose membrane and identified by specific antibodies followed by immunodetection performed using SuperSignal West Pico (34579) or Femto (34096) Plus Chemiluminescent Substrate (Thermo Fisher).

### Immunofluorescence analysis

HEK293T cells were cotransfected by the calcium phosphate method with pcDNA4-IRF2BP2 and pcDNA5-UBE2I to evaluate the immunolocalization. Forty-eight hours later, cells grown on glass coverslips were fixed with 4% paraformaldehyde. IRF2BP2 was detected by anti-c-Myc tag antibody, and UBE2I was detected by anti-UBE2I followed by Rhodamine- and Alexa 488-conjugated antibodies, respectively. For further assays, the cells were transfected with pcDNA4-IRF2BP2, pcDNA4-IRF2BP2-KR, or pcDNA5-SUMO1 as indicated. 4’,6’-diamidino-2-phenylindole (DAPI) staining was used to stain DNA and to indicate the nucleus position. The cells were observed, and images acquired at Confocal or Zeiss microscopy, as indicated in the figures.

## ACKNOWLEDGEMENTS

We are grateful to Leonardo Karan Teixeira (INCA) and Claudio Akio Masuda (UFRJ) for all suggestions and comments on the work. We thank Andres Cardozo Gizzi for the pmCherry and psYFP plasmids kindly donated, Gabriela Nestal for the INI-43 and Ivermectin inhibitors kindly gifted, and Marcelo Alex de Carvalho for single-lysine ubiquitin mutants kindly gifted. This work was supported by grants to J.P.B.V. from CNPq (311926/2021-5), FAPERJ (200.568/2023 and 200.435/2026). R.R.-O. and V.T.S. were supported by Brazilian Ministry of Health fellowship.

## AUTHOR CONTRIBUTIONS

R.R.-O conceived, designed, performed the experiments, analyzed the data and wrote the paper. V.T.S constructed the FRET plasmids. M.A.R. performed the FRET experiments and analyzed it. F.R.G.C. performed experiments and wrote the paper. J.P.B.V. designed experiments, analyzed the data and wrote the paper.

## REFERENCES

1. Childs, K. S., and Goodbourn, S. (2003) Identification of novel co-repressor molecules for Interferon Regulatory Factor-2 Nucleic Acids Res 31, 3016–3026

2. Yeung, K. T., Das, S., Zhang, J., Lomniczi, A., Ojeda, S. R., Xu, C. F. et al. (2011) A novel transcription complex that selectively modulates apoptosis of breast cancer cells through regulation of FASTKD2 Mol Cell Biol 31, 2287–2298

3. Urnov, F. D. (2002) A feel for the template: zinc finger protein transcription factors and chromatin Biochem Cell Biol 80, 321–333

4. Stadhouders, R., Cico, A., Stephen, T., Thongjuea, S., Kolovos, P., Baymaz, H. I., et al. (2015) Control of developmentally primed erythroid genes by combinatorial co-repressor actions Nat Commun 6, 8893

5. Wang, L., Gao, S., Wang, H., Xue, C., Liu, X., Yuan, H. et al. (2020) Interferon regulatory factor 2 binding protein 2b regulates neutrophil versus macrophage fate during zebrafish definitive myelopoiesis Haematologica 105, 325–337

6. Carneiro, F. R., Ramalho-Oliveira, R., Mognol, G. P., and Viola, J. P. (2011) Interferon regulatory factor 2 binding protein 2 is a new NFAT1 partner and represses its transcriptional activity Mol Cell Biol 31, 2889–2901

7. Ramalho-Oliveira, R., Oliveira-Vieira, B., and Viola, J. P. B. (2019) IRF2BP2: A new player in the regulation of cell homeostasis J Leukoc Biol 106, 717–723

8. Chen, H. H., Keyhanian, K., Zhou, X., Vilmundarson, R. O., Almontashiri, N. A., Cruz, S. A. et al. (2015) IRF2BP2 Reduces Macrophage Inflammation and Susceptibility to Atherosclerosis Circ Res 117, 671–683

9. Koeppel, M., van Heeringen, S. J., Smeenk, L., Navis, A. C., Janssen-Megens, E. M., and Lohrum, M. (2009) The novel p53 target gene IRF2BP2 participates in cell survival during the p53 stress response Nucleic Acids Res 37, 322–335

10. Teng, A. C., Kuraitis, D., Deeke, S. A., Ahmadi, A., Dugan, S. G., Cheng, B. L. et al. (2010) IRF2BP2 is a skeletal and cardiac muscle-enriched ischemia-inducible activator of VEGFA expression Faseb j 24, 4825–4834

11. Wang, G., Lu, T., Zhang, L., and Ding, J. (2024) IRF2BP2 binds to a conserved RxSVI motif of protein partners and regulates megakaryocytic differentiation Nature Communications 15, 10425

12. Sécca, C., Faget, D. V., Hanschke, S. C., Carneiro, M. S., Bonamino, M. H., de-Araujo-Souza, P. S., et al. (2016) IRF2BP2 transcriptional repressor restrains naive CD4 T cell activation and clonal expansion induced by TCR triggering J Leukoc Biol 100, 1081–1091

13. Palmroth, M., Viskari, H., Seppänen, M. R. J., Keskitalo, S., Virtanen, A., Varjosalo, M. et al. (2021) IRF2BP2 Mutation Is Associated with Increased STAT1 and STAT5 Activation in Two Family Members with Inflammatory Conditions and Lymphopenia Pharmaceuticals (Basel) 14,

14. García-Aznar, J. M., Maneiro Pampín, E., García Ramos, M., Acuña Pérez, M. J., Paz Gandiaga, N., Minguell Domingo, L. et al. (2023) Novel frameshift variants expand the map of the genetic defects in IRF2BP2 Front Immunol 14, 1279171

15. Fischer, S., Weber, L. M., Stielow, B., Frech, M., Simon, C., Geller, M. et al. (2024) IRF2BP2 counteracts the ATF7/JDP2 AP-1 heterodimer to prevent inflammatory overactivation in acute myeloid leukemia (AML) cells Nucleic Acids Res 52, 7590–7609

16. Endo, Y., Koga, T., Kurushima, S., Tomokawa, T., Mishima, H., Yoshiura, K.-I., et al. (2025) Case Report: Novel IRF2BP2 variant in a Japanese patient with impaired B-cell differentiation, Th1 polarization, and systemic immune dysregulation Frontiers in Immunology **Volume** 16 - 2025,

17. Cruz, S. A., Hari, A., Qin, Z., Couture, P., Huang, H., Lagace, D. C., et al. (2017) Loss of IRF2BP2 in Microglia Increases Inflammation and Functional Deficits after Focal Ischemic Brain Injury Front Cell Neurosci 11, 201

18. Feng, X., Lu, T., Li, J., Yang, R., Hu, L., Ye, Y. et al. (2020) The Tumor Suppressor Interferon Regulatory Factor 2 Binding Protein 2 Regulates Hippo Pathway in Liver Cancer by a Feedback Loop in Mice Hepatology 71, 1988–2004

19. Yao, Y., Wang, Y., Li, L., Xiang, X., Li, J., Chen, J. et al. (2019) Down-regulation of interferon regulatory factor 2 binding protein 2 suppresses gastric cancer progression by negatively regulating connective tissue growth factor J Cell Mol Med 23, 8076–8089

20. He, J., McLaughlin, R. P., van der Beek, L., Canisius, S., Wessels, L., Smid, M. et al. (2020) Integrative analysis of genomic amplification-dependent expression and loss-of-function screen identifies ASAP1 as a driver gene in triple-negative breast cancer progression Oncogene 39, 4118–4131

21. Pastor, T. P., Peixoto, B. C., and Viola, J. P. B. (2021) The Transcriptional Co-factor IRF2BP2: A New Player in Tumor Development and Microenvironment Frontiers in Cell and Developmental Biology Volume 9 - 2021,

22. Chen, Y., Zhuo, R., Sun, L., Tao, Y., Li, G., Zhu, F. et al. (2024) Super-enhancer-driven IRF2BP2 enhances ALK activity and promotes neuroblastoma cell proliferation Neuro Oncol 26, 1878–1894

23. Yu, J., Zhang, Z., Chen, Y., Wang, J., Li, G., Tao, Y., et al. (2025) Super-Enhancer-Driven IRF2BP2 is Activated by Master Transcription Factors and Sustains T-ALL Cell Growth and Survival Adv Sci (Weinh) 12, e2407113

24. Fang, J., Ji, Y. X., Zhang, P., Cheng, L., Chen, Y., Chen, J. et al. (2020) Hepatic IRF2BP2 Mitigates Nonalcoholic Fatty Liver Disease by Directly Repressing the Transcription of ATF3 Hepatology 71, 1592–1608

25. Li, T., Luo, Q., He, L., Li, D., Li, Q., Wang, C. et al. (2020) Interferon Regulatory Factor-2 Binding Protein 2 Ameliorates Sepsis-Induced Cardiomyopathy via AMPK-Mediated Anti-Inflammation and Anti-Apoptosis Inflammation 43, 1464–1475

26. Ellegast, J. M., Alexe, G., Hamze, A., Lin, S., Uckelmann, H. J., Rauch, P. J. et al. (2022) Unleashing Cell-Intrinsic Inflammation as a Strategy to Kill AML Blasts Cancer Discov 12, 1760–1781

27. Schuh, K., Twardzik, T., Kneitz, B., Heyer, J., Schimpl, A., and Serfling, E. (1998) The interleukin 2 receptor alpha chain/CD25 promoter is a target for nuclear factor of activated T cells J Exp Med 188, 1369–1373

28. Pan, Y., Shang, G., Li, J., Zhang, Y., Liu, J., Ji, Y., et al. (2023) Case Report: A novel IRF2BP2 mutation in an IEI patient with recurrent infections and autoimmune disorders Front Immunol 14, 967345

29. Soliman, H., Khalil, F., and Antonia, S. (2014) PD-L1 expression is increased in a subset of basal type breast cancer cells PLoS One 9, e88557

30. Dorand, R. D., Nthale, J., Myers, J. T., Barkauskas, D. S., Avril, S., Chirieleison, S. M. et al. (2016) Cdk5 disruption attenuates tumor PD-L1 expression and promotes antitumor immunity Science 353, 399–403

31. Wu, A., Wu, Q., Deng, Y., Liu, Y., Lu, J., Liu, L., et al. (2019) Loss of VGLL4 suppresses tumor PD-L1 expression and immune evasion Embo j 38,

32. Cui, Y., Zhao, Y., Shen, G., Lv, Q., and Ma, L. (2024) CDYL loss promotes cervical cancer aggression by increasing PD-L1 expression via the suppression of IRF2BP2 transcription Transl Oncol 47, 102038

33. Nyquist, K. B., Panagopoulos, I., Thorsen, J., Haugom, L., Gorunova, L., Bjerkehagen, B. et al. (2012) Whole-transcriptome sequencing identifies novel IRF2BP2-CDX1 fusion gene brought about by translocation t(1;5)(q42;q32) in mesenchymal chondrosarcoma PLoS One 7, e49705

34. Panagopoulos, I., Gorunova, L., Lobmaier, I., and Heim, S. (2023) Fusion of the Genes for Interferon Regulatory Factor 2 Binding Protein 2 (IRF2BP2) and Caudal Type Homeobox 1 (CDX1) in a Chondrogenic Tumor In Vivo 37, 2459–2463

35. Shimomura, Y., Mitsui, H., Yamashita, Y., Kamae, T., Kanai, A., Matsui, H. et al. (2016) New variant of acute promyelocytic leukemia with IRF2BP2-RARA fusion Cancer Sci 107, 1165–1168

36. Arruda, L. C., Lorenzi, J. C., Sousa, A. P., Zanette, D. L., Palma, P. V., Panepucci, R. A. et al. (2015) Autologous hematopoietic SCT normalizes miR-16, -155 and -142-3p expression in multiple sclerosis patients Bone Marrow Transplant 50, 380–389

37. van Solingen, C., Cyr, Y., Scacalossi, K. R., de Vries, M., Barrett, T. J., de Jong, A., et al. (2022) Long noncoding RNA CHROMR regulates antiviral immunity in humans Proc Natl Acad Sci U S A 119, e2210321119

38. Barysch, S. V., Stankovic-Valentin, N., Miedema, T., Karaca, S., Doppel, J., Nait Achour, T., et al. (2021) Transient deSUMOylation of IRF2BP proteins controls early transcription in EGFR signaling EMBO Rep 22, e49651

39. Zanchin, N. I., and Goldfarb, D. S. (1999) Nip7p interacts with Nop8p, an essential nucleolar protein required for 60S ribosome biogenesis, and the exosome subunit Rrp43p Mol Cell Biol 19, 1518–1525

40. Karpova, T. S., Baumann, C. T., He, L., Wu, X., Grammer, A., Lipsky, P. et al. (2003) Fluorescence resonance energy transfer from cyan to yellow fluorescent protein detected by acceptor photobleaching using confocal microscopy and a single laser J Microsc 209, 56–70

41. Jares-Erijman, E. A., and Jovin, T. M. (2003) FRET imaging Nat Biotechnol 21, 1387–1395

42. Xue, Y., Zhou, F., Fu, C., Xu, Y., and Yao, X. (2006) SUMOsp: a web server for sumoylation site prediction Nucleic Acids Res 34, W254–257

43. Gong, L., Kamitani, T., Fujise, K., Caskey, L. S., and Yeh, E. T. (1997) Preferential interaction of sentrin with a ubiquitin-conjugating enzyme, Ubc9 J Biol Chem 272, 28198–28201

44. Giorgino, F., de Robertis, O., Laviola, L., Montrone, C., Perrini, S., McCowen, K. C. et al. (2000) The sentrin-conjugating enzyme mUbc9 interacts with GLUT4 and GLUT1 glucose transporters and regulates transporter levels in skeletal muscle cells Proc Natl Acad Sci U S A 97, 1125–1130

45. Gareau, J. R., and Lima, C. D. (2010) The SUMO pathway: emerging mechanisms that shape specificity, conjugation and recognition Nat Rev Mol Cell Biol 11, 861–871

46. El-Asmi, F., McManus, F. P., Thibault, P., and Chelbi-Alix, M. K. (2020) Interferon, restriction factors and SUMO pathways Cytokine Growth Factor Rev 55, 37–47

47. Melchior, F. (2000) SUMO--nonclassical ubiquitin Annu Rev Cell Dev Biol 16, 591–626

48. Teng, A. C., Al-Montashiri, N. A., Cheng, B. L., Lou, P., Ozmizrak, P., Chen, H. H., et al. (2011) Identification of a phosphorylation-dependent nuclear localization motif in interferon regulatory factor 2 binding protein 2 PLoS One 6, e24100

49. Christie, M., Chang, C. W., Róna, G., Smith, K. M., Stewart, A. G., Takeda, A. A. et al. (2016) Structural Biology and Regulation of Protein Import into the Nucleus J Mol Biol 428, 2060–2090

50. Lange, A., Mills, R. E., Lange, C. J., Stewart, M., Devine, S. E., and Corbett, A. H. (2007) Classical nuclear localization signals: definition, function, and interaction with importin alpha J Biol Chem 282, 5101–5105

51. Wagstaff, K. M., Sivakumaran, H., Heaton, S. M., Harrich, D., and Jans, D. A. (2012) Ivermectin is a specific inhibitor of importin α/β-mediated nuclear import able to inhibit replication of HIV-1 and dengue virus Biochem J 443, 851–856

52. van der Watt, P. J., Chi, A., Stelma, T., Stowell, C., Strydom, E., Carden, S. et al. (2016) Targeting the Nuclear Import Receptor Kpnβ1 as an Anticancer Therapeutic Mol Cancer Ther 15, 560–573

53. Lamsoul, I., Lodewick, J., Lebrun, S., Brasseur, R., Burny, A., Gaynor, R. B. et al. (2005) Exclusive ubiquitination and sumoylation on overlapping lysine residues mediate NF-kappaB activation by the human T-cell leukemia virus tax oncoprotein Mol Cell Biol 25, 10391–10406

54. Liebelt, F., and Vertegaal, A. C. (2016) Ubiquitin-dependent and independent roles of SUMO in proteostasis Am J Physiol Cell Physiol 311, C284–296

55. Glickman, M. H., and Ciechanover, A. (2002) The Ubiquitin-Proteasome Proteolytic Pathway: Destruction for the Sake of Construction Physiological Reviews 82, 373–428

56. Sun, M., and Zhang, X. (2022) Current methodologies in protein ubiquitination characterization: from ubiquitinated protein to ubiquitin chain architecture Cell Biosci 12, 126

57. Michel, M. A., Swatek, K. N., Hospenthal, M. K., and Komander, D. (2017) Ubiquitin Linkage-Specific Affimers Reveal Insights into K6-Linked Ubiquitin Signaling Mol Cell 68, 233–246.e235

58. Flotho, A., and Melchior, F. (2013) Sumoylation: a regulatory protein modification in health and disease Annu Rev Biochem 82, 357–385

59. Han, Z. J., Feng, Y. H., Gu, B. H., Li, Y. M., and Chen, H. (2018) The post-translational modification, SUMOylation, and cancer (Review) Int J Oncol 52, 1081–1094

60. Zhou, Z., Wang, M., Li, J., Xiao, M., Chin, Y. E., Cheng, J. et al. (2016) SUMOylation and SENP3 regulate STAT3 activation in head and neck cancer Oncogene 35, 5826–5838

61. Hsu, C. Y., Yeh, L. T., Fu, S. H., Chien, M. W., Liu, Y. W., Miaw, S. C. et al. (2018) SUMO-defective c-Maf preferentially transactivates Il21 to exacerbate autoimmune diabetes J Clin Invest 128, 3779–3793

62. Wilkinson, K. A., Nakamura, Y., and Henley, J. M. (2010) Targets and consequences of protein SUMOylation in neurons Brain Res Rev 64, 195–212

63. Ihara, M., Stein, P., and Schultz, R. M. (2008) UBE2I (UBC9), a SUMO-conjugating enzyme, localizes to nuclear speckles and stimulates transcription in mouse oocytes Biol Reprod 79, 906–913

64. Li, F., Lai, L., You, Z., Cheng, H., Guo, G., Tang, C., et al. (2022) Identification of UBE2I as a Novel Biomarker in ccRCC Based on a Large-Scale CRISPR-Cas9 Screening Database and Immunohistochemistry Frontiers in Molecular Biosciences Volume 9 - 2022,

65. Xu, H., and Xu, B. (2024) UBE2I regulates the nuclear translocation of hnRNPA2B1 by contributing to SUMO modification in osteoarthritis Gene 927, 148740

66. Bartel, P. L., and Fields, S. (1995) Analyzing protein-protein interactions using two-hybrid system Methods Enzymol 254, 241–263

67. Otto, T. D., Vasconcellos, E. A., Gomes, L. H. F., Moreira, A. S., Degrave, W. M., Mendonça-Lima, L. et al. ChromaPipe: a pipeline for analysis, quality control and management for a DNA sequencing facility

68. Hara, K., Maruki, Y., Long, X., Yoshino, K., Oshiro, N., Hidayat, S. et al. (2002) Raptor, a binding partner of target of rapamycin (TOR), mediates TOR action Cell 110, 177–189

