## SUPPORTING INFORMATION for "SUMOylation Regulates IRF2BP2 Nuclear Localization and Stability"

**Contents:** Figure S1, Tables S1-S3.

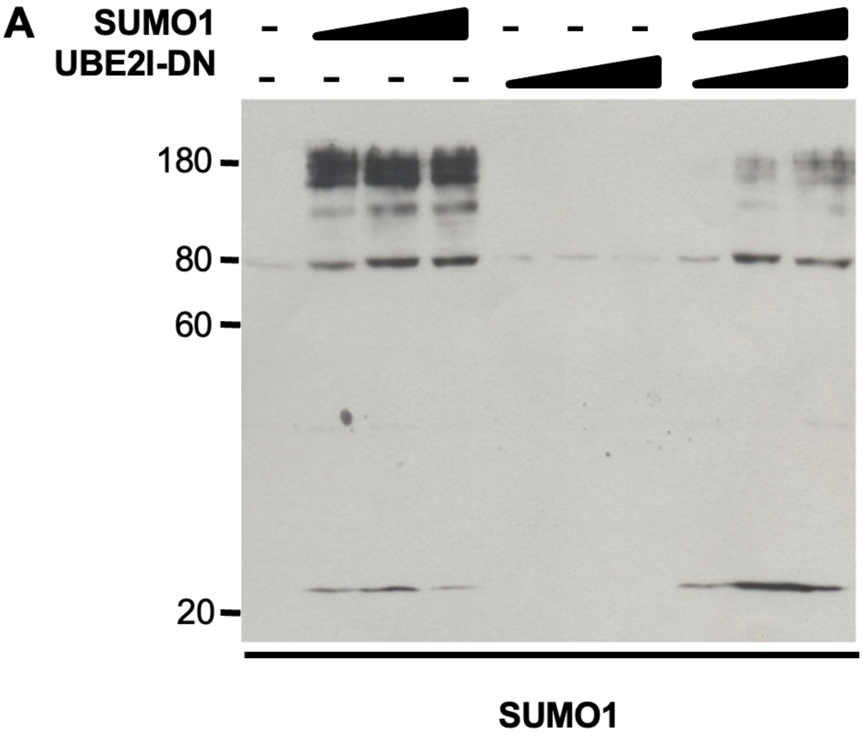

**Figure S1. Dominant-negative UBE2I block SUMO conjugation.** Total protein extracts of HEK293T cells expressing increasing amounts of SUMO1 and/or UBE2I-DN were obtained. Equal quantities of protein were fractioned by SDS-PAGE and subjected to immunoblot using anti-SUMO1 antibody.

**TABLE S1. Plasmids.**

| Plasmid | 5' site | 3' site | Sequence (nt) | Protein | Vector backbone |
|---|---|---|---|---|---|
| pTL-IRF2BP2 (NM_001077397.1) | EcoRI | SalI | 1-1716 | LexA(DBD)-IRF2BP2 | pTL-1 Carneiro et al., 2006 |
| pTL-IRF2BP2-RING | EcoRI | SalI | 903-1716 | LexA(DBD)-IRF2BP2-RING | pTL-1 Carneiro et al., 2006 |
| pTL-IRF2BP2-ZINC | EcoRI | XhoI | 1-1452 | LexA(DBD)-IRF2BP2-ZINC | pTL-1 Carneiro et al., 2006 |
| pmCHERRY-IRF2BP2 | BglII | HindIII | 1-1713 | mCherry-IRF2BP2 | pmCherry-N1 |
| psYFP-UBE2I (NM_003345.3) | BglII | HindIII | 1-474 | sYFP-UBE2I | psYFP-N1 |
| pcDNA4-IRF2BP2 Carneiro *et al*., 2011 | EcoRI | XhoI | 1-1713 | IRF2BP2-c-myc-His (IRF2BP2) | pcDNA4-To/myc-HisA |
| pcDNA5-UBE2I (NM_003345.3) | BamHI | XhoI | 1-477 | UBE2I | pcDNA5-FRT0 |
| pcDNA5-SUMO1 (NM_003352.4) | BamHI | XhoI | 1-306 | SUMO1 | pcDNA5-FRT0 |
| pcDNA4-IRF2BP2-AAAA (MutNLS) | Carneiro *et al*., 2011 | | | IRF2BP2-AAAA | pcDNA4-To/myc-HisA |
| pcDNA4-IRF2BP2-KR | Lysine residues 566, 326, 82, and 372 were mutated to Arginine by Genescript | | | IRF2BP2-c-myc-His-KR (IRF2BP2-KR) | pcDNA4-To/myc-HisA |
| pTL-Nip7 | Zanchin and Goldfarb, 1999 | | | lexA(DBD)-Nip7 | pTL-1 Carneiro et al., 2006 |
| pACT-Nop8 | Zanchin and Goldfarb, 1999 | | | LexA(AD)-Nop8 | |

**TABLE S2. Primers used in the plasmid constructions**

| Plasmid | Foward | Reverse |
|---|---|---|
| pTL-IRF2BP2 | 5’GATC **GAA TTC** ATG GCC GCG GCG GTG GCG 3’ | 5’ CCTA **GTC GAC** TTA CGA GTC TCT CTC TTT TTT C 3’ |
| pTL-IRF2BP2-RING | 5’ GATC **GAA TTC** CCC AAG ACC GTG CGC GAC 3’ | 5’ CCTA **GTC GAC** TTA CGA GTC TCT CTC TTT TTT C 3’ |
| pTL-IRF2BP2-ZINC | 5’GATC **GAA TTC** ATG GCC GCG GCG GTG GCG 3’ | 5’ CGTA **CTC GAG** TTA ACT GGT TGC CAG AGA GGA GTC 3’ |
| pmCHERRY-IRF2BP2 | AGTCAGATCTATATGGCCGCGGCGGTGGCGGTG | AGGCAAGCTTCGAGTCTCTCTCTTTTTTC |
| psYFP-UBE2I | AGTCAGATCTATGTCGGGGATCGCCCTC | AGTCAAGCTTTGAGGGCGCAAACTTC |
| pcDNA4-IRF2BP2 | 5’ CATG **GAA TTC** GGC TCC TCG GAC ATG GCC G 3’ | 5’ CCTA **CTC GAG** CGA GTC TCT CTC TTT TTT CAC 3’ |
| pcDNA5-UBE2I | 5' GAT C**GG ATC C**AC TTT GAA CAT GTC GGG GAT CG 3' | 5' GAT C**CT CGA G**TG AGG GCG CAA ACT TC 3' |
| pcDNA5- | 5' CTT CGG GGA CAG TGT CCC TGT CCA TCT TAG A 3' | 5' TCT AAG ATG GAC AGG GAC ACT GTC CCC GAA G 3' |
| pcDNA5-SUMO1 | 5' GAT C**GG ATC C**AT GTC TGA CCA GGA 3' | 5' GAT CCT CGA GAA CTG TTG AAT GAC CCC CC 3' |

In bold: restriction sites

**TABLE S3. Antibodies and drugs**

| **Antibody** | **SOURCE** | **IDENTIFIER** | **DILUTION/DOSE** |
|---|---|---|---|
| IRF2BP2 | Santa Cruz | sc-136715 | 1:500 |
| c-Myc-Tag | Invitrogen | Invitrogen 46-0306 | 1:2500; 2µg for immunoprecipitation |
| UBE2I | Santa Cruz | sc-10759 | 1:1000 |
| SUMO1 | Santa Cruz | sc-5308 | 1:1000 |
| GAPDH | Santa Cruz | sc-32233 | 1:2500 |
| HA | Santa Cruz | Sc-7392 | 1:1000 |
| Histone H3 | Cell Signaling | 9712 | 1:1000 |
| Histone H3-K4me3 | Cell Signaling | 9727 | 1:1000 |
| Histone H3-K27me3 | Cell Signaling | 9733 | 1:1600 |
| Alexa 488 | Thermo | A32723TR | 1:1000 |
| Alexa 594 conjugated antibody | Thermo | 31660 | 1:1000 |
| **Analogs** | | | |
| BrDU | Phoenix Flow Systems | ABFM18 | 20 µM for 30 minutes |
| **Inhibitors** | | | |
| INI43 | Sigma | SML1911 | 10 µM for 1 hour |
| Ivermectin | Sigma | I8898 | 10 µM for 1 hour |
| MG132 | Sigma | 474790 | 20 µM for 2h or 24h, as indicated in the figure |
| Thymidine | Merk Millipore | 6060 | 2.5 mM 18h |
